# Identifying Protein Complexes in Protein-protein Interaction Data using Graph Convolution Network

**DOI:** 10.1101/2021.07.07.451457

**Authors:** Nazar Zaki, Harsh Singh

## Abstract

Protein complexes are groups of two or more polypeptide chains that join together to build noncovalent networks of protein interactions. A number of means of computing the ways in which protein complexes and their members can be identified from these interaction networks have been created. While most of the existing methods identify protein complexes from the protein-protein interaction networks (PPIs) at a fairly decent level, the applicability of advanced graph network methods has not yet been adequately investigated. In this paper, we proposed various graph convolutional networks (GCNs) methods to improve the detection of the protein functional complexes. We first formulated the protein complex detection problem as a node classification problem. Second, the Neural Overlapping Community Detection (NOCD) model was applied to cluster the nodes (proteins) using a complex affiliation matrix. A representation learning approach, which combines the multi-class GCN feature extractor (to obtain the features of the nodes) and the mean shift clustering algorithm (to perform clustering), is also presented. We have also improved the efficiency of the multi-class GCN network to reduce space and time complexities by converting the dense-dense matrix operations into dense-spares or sparse-sparse matrix operations. This proposed solution significantly improves the scalability of the existing GCN network. Finally, we apply clustering aggregation to find the best protein complexes. A grid search was performed on various detected complexes obtained by applying three well-known protein detection methods namely ClusterONE, CMC, and PEWCC with the help of the Meta-Clustering Algorithm (MCLA) and Hybrid Bipartite Graph Formulation (HBGF) algorithm. The proposed GCN-based methods were tested on various publicly available datasets and provided significantly better performance than the previous state-of-the-art methods. The code and data used in this study are available from https://github.com/Analystharsh/GCN_complex_detection

## 1. Introduction

Proteins are responsible for the growth and development of all species. The majority of the functions in the cellular systems of living beings are not caused by the individual protein nodes. Instead, many protein nodes take part in performing cellular functions. These similar protein nodes are known as protein complexes or protein communities. The overwhelming majority of biological processes within cells are controlled by proteins; thus, proteins control appropriate cell functionality. Cells are required to respond to several stimuli. The cellular response is a complex procedure involving the assignation of particular tasks to specific proteins; for any function, a specific type and number of proteins will be needed. For this reason, biologists have moved from examining the relationships between structure and function in individual protein families to making intensive studies of complete cellular networks [1]. For complete comprehension of the function of a protein, it must be examined in the light of those partners with which it interacts and the complex to which it belongs. It is generally recognized that protein complexes comprise groups of two or more interacting polypeptide chains [2]. Being able to detect such complexes has a significant impact as they are central to biological processes and create the framework for the network of protein-protein interactions (PPIs). Protein complex formation is responsible for antigen-antibody interaction and transportation, gene expression control, cell cycle control, signaling, differentiation, protein folding, transcription, translation, and inhibition of enzymes [2]. Thus, building or destroying various protein complexes causes the initialization, modulation, or termination of several biological processes [3]. Gene mutations can cause a considerable quantity of protein complex abnormalities [4], [5]. Subsequently, this may influence how proteins interact with other partners. In particular, it may modify the ways in which different proteins interact, and in certain instances, it may also initiate self-interaction [4]. These modifications are small but have significance as they are allied to significant numbers of alterations in self-functionality, which can assist us to achieve a better understanding of ways to engender a resolution. Additionally, knowledge of protein complexes can increase knowledge of various forms of the disease. Much research has demonstrated that genetic disease is caused by proteins with similar functional interactions [2], [6]. Using data extracted from PPI assist researchers in the discovery of inter-gene evolutionary relationships that can guide them to unique protein complexes, leading to the discovery of unique genes related to particular diseases [4]. Additionally, protein complexes are making significant changes in terms of the creation of new drug therapies [7]. This is because they play a significant part in physiological function, which makes them superior to standard in vitro methods of analyzing therapeutic agents [7]. The investigation of protein complexes may reveal previously undiscovered pathways and proteins, new methods for controlling diseases, and new ways of classifying genes. All of these elements combine to allow for the development and understanding of the targeting, identification, and retardation of disease progression.

Many previous successful methods have been put forward for the detection of protein complexes from PPI networks which can be divided into the following seven categories:

1. A local neighborhood density search approach, focusing on the discovery of dense subgraphs inside the input network, including MCODE [8], DPClus [7], ProRank [9], [10], ProRank+ [11], CMC [12], PROCOMOSS [13] and PEWCC [14], NCMine [15], Core&Peel [16], SPICi [17] and non-cooperative sequential game [18].
2. Local search approaches based on cost, focusing on the extraction of modules from interaction graphs through the partition of the graphs into linked subgraphs employing cost functions for the guidance of searches towards the optimal partition, including RNSC [19], ModuLand [20], and STM [21].
3. Approaches employing Flow Simulation, focusing on the imitation of ways in which information spreads through a network, including MCL [22] and RW [6].
4. Approaches based on statistics, relying on the employment of statistical concepts for clustering proteins, e.g. how many shared neighbors pair of proteins have, and on notions of referential attachments for module members with other elements within the module; this includes SL [23], idenPC-MIIP [24], idenPC-CAP [25] and Farutin [2].
5. Stochastic search methods based on population employed to develop algorithms used to detect communities and networks including CGA [26], IGA [7], and EHO-MCL [27].
6. Approaches based on modularity, topological structure, overlapping information, and GO annotations, including CFinder [28] and [29], ClusterONE [30], and SE-DMTG [31].
7. Graph-based clustering methods, which includes statistical-based measures methods such as [2] which uses the concept of statistical significance to measure the strength of the relationship between two nodes (proteins), which requires prior estimation of the p-value. Cost-based Local search (CL) [32], Population-based Stochastic search (PS) [26] and [33], Local neighborhood Density search (LD), [34] and [30].

While all of the methods detailed above can identify protein complexes at a fairly accurate level, in this paper, we introduce four major contributions to improve the detection of the protein complexes from the PPI network. These contributions are as follow:

- **Contribution 1:** we employ the node classification approaches [35] to classify nodes (proteins) into classes (complexes). First, the interaction matrix (adjacency matrix) and degree matrix are prepared from a given PPI network. Second, the identity matrix is used as a feature of the nodes. Based on all these inputs, three versions of GCN [36] were employed. These models are multi-class GCN classification methods with 2^*N*^ label size (*N* is the number of complexes), multi-class GCN classification with label size *K* (*K* is the number of possible combinations of the labels in the respective datasets), and multi-label GCN classification method. All these classification methods provided the protein complex labels for all the nodes in the network. These models not only detect non-overlapping communities but also are self-sufficient in the definition itself to detect overlapping complexes [37].
- **Contribution 2:** GCN approaches are further advanced by proposing efficient matrix operations inside GCN layers. It leads to the decrement in the time and space complexities required to implement, train, and infer the GCN approaches conveniently. The dense matrices involved in the GCN model, such as feature, adjacency, and degree matrices are turned into the compressed sparse row (CSR) matrix format [38]. It removes the redundant operations from the existing GCN architectures.
- **Contribution 3:** Two learning representations were proposed for complex detection. The first approach NOCD GCN method [39] uses the generative model to learn the community affiliation matrix from the node features and adjacency matrix. It requires modeling the loss function in terms of negative likelihood involving the Bernoulli Poisson (BP) method. The second approach extracts feature from the existing pre-trained GCN model and use these feature embedding to form clustering with mean shift algorithm [40].
- **Contribution 4:** A clustering aggregation process [41] has been proposed, which takes all the clustering from different methods without knowing which one is the best and produces the optimal clustering. A grid search has been performed with the Meta-Clustering Algorithm [42] and Hybrid Bipartite graph formulation (HBGF) [43] algorithm.

## 2. Methods

### 2.1. Datasets

#### 2.1.1. Human and Mouse datasets

The interaction networks of these datasets are extracted from the BioGrid interaction database [44]. These datasets provide PPI networks of two species “Homo sapiens” and “Mus musculus” popularly known as human and mouse, respectively. These raw datasets are pre-processed by removing duplicate nodes/interaction edges and merging all the available PPIs for particular species. The Human PPI network contains 21,644 protein nodes and 338,923 interactions, while the Mouse PPI contains 21,420 protein nodes and 20,679 interactions. The CORUM reference complexes dataset [45] which includes 623 “Mouse” related reference complexes and 2,645 “human” related complexes were used to evaluate the performance of the proposed methods.

#### 2.1.2. Yeast PPI datasets

In addition, two popular datasets Collins and Gavin datasets [46] were also considered in this study. Gavin dataset is extracted by calculating socio-affinity index in all yeast PPI networks from the database, as proposed by original authors. If this term is greater than 5, then that PPI is considered. If not, then it is not included. On the other side, a different metric (purification enrichment test) is used to retain the Collins dataset.

### 2.2. Developing the multi-class GCN classification model

Inspired by the recent promising results achieved by applying graph-based learning techniques for detecting communities in graphs [47], we employ the GCN [48] to classify proteins into complexes. The method starts by creating an adjacency matrix *Â* of shape *N* × *N* (where *N* is the number of nodes. In this case, if two nodes are connected in the input graph, then the corresponding entry in the adjacency matrix is presented as 1 or 0 otherwise. Therefore, the input feature matrix *F*_1_ is considered as an identity matrix of shape *N* × *N*, which presents the features for each node in the absence of explicit node features. The input adjacency matrix is formed by adding the feature matrix *F*_1_ and *Â* to add self-connection of each node in the adjacency matrix (*A* = *Â* + *F*_1_). This input feature matrix is then normalized [36] as *Z* = *D*^−1/2^*A D*^−1/2^. In this case, each graph neural network layer will consist of its weight matrices *W*_*i*_. The *W*_1_for example is the weight matrix of shape *N* × 512 for the first GCN layer [48]. We can then obtain the matrix *K* by multiplying the feature matrix *F*_1_ by weight matrix *W*_1_ (*K* = *F*_1_*W*_1_). Further, the matrix *K* is multiplied by *Z* to obtain *P* (*P* = *ZK*). Which then passed to the ReLU activation function *F*_2_ = *ReLU*(*P*). This output is again passed to another GCN layer with a weight matrix *W*_2_ of shape 512 × *L* as a feature matrix for the next layer, where *L* is the length of each node label. This process is repeated using *F*_2_, *W*_2_, *A*, and *D*.

The label matrix *Y* for all the nodes in the input graph is prepared by inspecting the possible number of combinations *K* for nodes belonging to one or more complexes. In this way, each label is formed as one hot encoder of the length equal to the number of such possible combinations for the particular dataset. Finally, the output *F*_3_ obtained from the second GCN layer is passed to the row-wise softmax function *Ŷ* = *σ*(*F*_3_) where *σ* is row-wise Softmax function. With the help of *Y* and *Ŷ*, the presented GCN is trained with the loss function defined as categorical cross-entropy *L* = *H*(*Y, Ŷ*), where *H*(. : .) represents categorical cross-entropy, *Y* and *Ŷ* are the ground truth label matrix and predicted outcome matrix, respectively. The weighted matrices *W*_1_ and *W*_2_ are updated during the training using multiple epochs.

To extend the method to handle multi-label classification, instead of the softmax function, we used the sigmoid function after the second GCN layer, and the loss function was also changed into a class-wise summation of binary cross-entropy. Therefore, the loss function will be 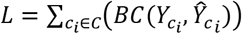 where *C* is the set of all classes, and *c*_*i*_ is class *i* of the set *C* and BC is the binary cross-entropy. During the training time, this loss function was used to calculate the gradient, while Adam optimizer was used to update the weights *W*_1_ and *W*_2_. Finally, the values > 0.5 are converted into 1, otherwise converted into 0. These steps are illustrated in Figure 1.

**Figure 1:**
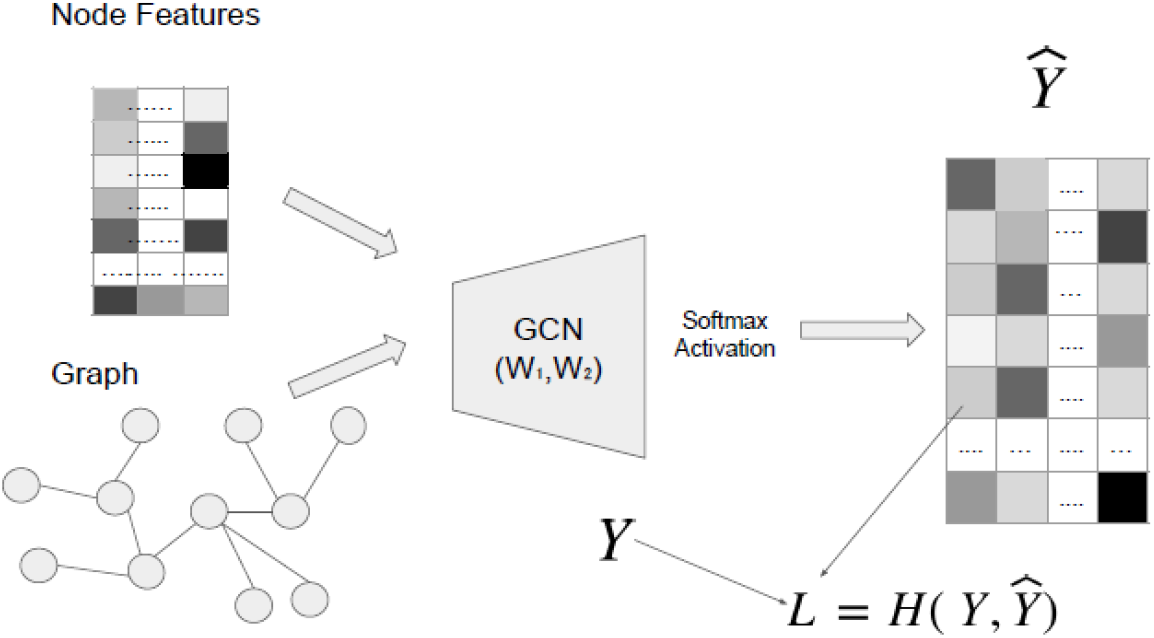
Overview of the multi-class GCN node classification

#### 2.2.1. Explicit node features

In our experiments, we have primarily used Identity matrix as feature embedding. Apart from this, we have also tried a custom feature matrix. The RNA-RNA interaction networks and RNA-protein interaction networks [49] were used to build the feature matrix. For the node classification task, the adjacency matrix and feature matrix are required as input. The process of obtaining the feature matrix (feature embedding) from the mentioned two extra networks is as follows:

- The RNA-RNA interaction networks are clustered using greedy modularity [50]. In this case, almost all the RNA nodes can be clustered into three big groups. Therefore, the feature length for every node is assigned to 3. It is one hot encoder with a length of 3. Entry 1 in this feature embedding represents the inclusion of the relationship with corresponding RNA clusters, while entry 0 represents the non-inclusion of that RNA-RNA cluster. The Girvan-Newman [51] yielded one big cluster (∼ 55600 nodes out of 56000 nodes in total)
- The RNA-RNA clusters are labeled with unique IDs,
- Every protein was labeled as per the RNA clusters from which they are connected in the RNA-Protein network. Generally, it is in sparse matrix form as most of the entries of this feature matrix are zeros.
- Use these node feature embedding as a feature matrix.

### 2.3. Representation Learning Approaches

One of the limitations of GCN manipulations is the computational cost. The process includes large and sparse matrix addition, matrix inversion, matrix multiplication, etc., and therefore, the use of dense matrices causes high computational costs. To address this issue we first represented the sparse matrix using the compressed sparse row (CSR) matrix format [52].

#### 2.3.1. The Neural Overlapping Community Detection - GCN method

In this section, we describe the probabilistic generative GCN model. A model which does not require any label for the training purpose. It simply takes the graph adjacency matrix and node feature matrix as inputs. Then, it uses Neural Overlapping Community Detection (NOCD) model [53] to learn the connectivity among the nodes (protein) and optimize the weights accordingly. It is a completely unsupervised method that relies on the probabilistic modeling of the output. This output is termed as complex affiliation matrix F. So the problem boils down to the *p*(*A*|*F*) estimation, where A is the adjacency matrix. If the complex affiliation matrix is prior, entries in the adjacency matrix can be sampled as, 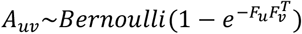, where *u* and *v* are two nodes. In this case, the higher value of the 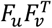 indicates the higher chances of *u* and *v* are connected and be in the same community. All the settings are kept the same as the multi-class GCN classification method, but the row-wise softmax function is changed to the element-wise Relu activation after the second GCN layer. The other two differences in the GCN architectures are as follow:

- The dropout layer is used in the last GCN layer
- The *L*_2_ regularization is applied to both weight matrices in the network
- The batch normalization layer is also added to the first GCN layer

In this case, the output here is complex affiliation matrix F, not *Ŷ*, which uses labels to perform the training.

Therefore, the negative likelihood of the proposed model can be written as:

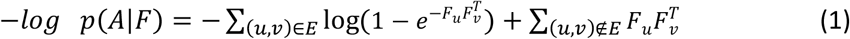

To reduce the effect of non-edges (sparse effect), we selected only a certain number of non-edges to balance the estimation. This new term is written as follows:

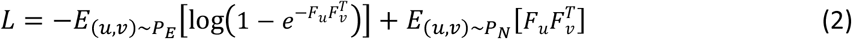

Where *P*_*E*_ and *P*_*N*_ indicate the uniform distribution over edges and non-edges. By minimizing this loss function, we can optimize the weights of the hidden layer of the GCN. The hidden layer has a size of 512 (Figure 2).

**Figure 2:**
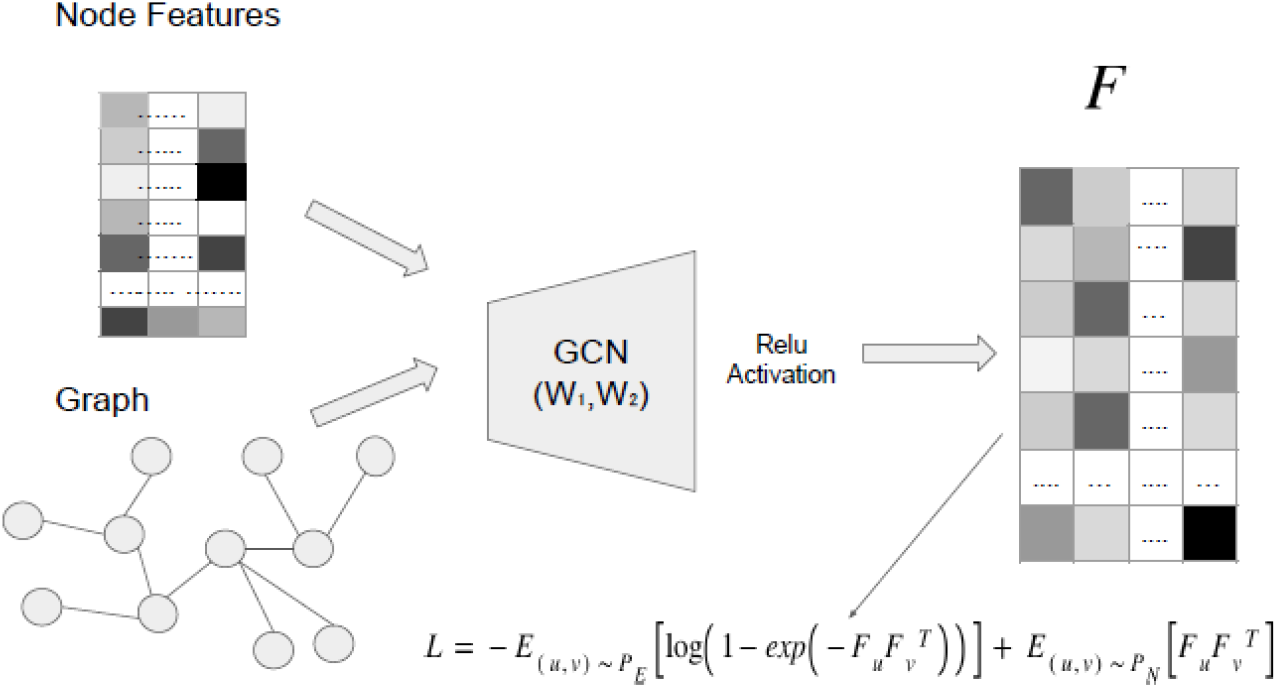
Overview of the NOCD GCN model.

#### 2.3.2. GCN feature extraction and unsupervised feature learning

In this step, we extracted features for all the nodes in the datasets from the last layer of the multi-class GCN classification model (before applying the row-wise softmax function) pre-trained on the Human and Mouse datasets.

The length of each of the feature vectors is equals to the number of complexes present in that particular dataset. Once these features are retrieved, we applied several unsupervised algorithms, which do not require prior information about the number of clusters.

One of such algorithms is the mean shift algorithm. This algorithm accepts the node features, estimates the number of clusters, and assigns nodes to clusters. The kernel that was used in this experimental work is the flat/uniform kernel. In this case, the kernel 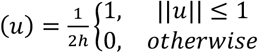, where *u* is the data point and *h* is the bandwidth of the kernel. This algorithm identifies dense regions by using the following kernel density estimation (KDE) function:

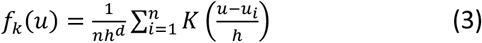

Where *n* is the total number of data points, and *u*_*i*_ is an *i*^*th*^ data point. First, this algorithm finds dense regions with predefined bandwidth. Second, it determines the mean of the data points in that region. It shifts its centroid toward the mean point. The second step is repeated until the shifting of centroids stops. It is also called the convergence of the mean shift algorithm. In this way, it computes the optimal number of clusters.

#### 2.3.3. Meta-Clustering Algorithm (MCLA)

The objective of this algorithm is to combine clusters obtained from different clustering techniques. It also provides the association confidence estimation of all the instances (or data points). MCLA uses hyperedges [54] as the starting vertices. In this case, the hyperedges are the members of the indicator matrix (consider indicator matrix as a set of column vectors) *H*^*l*^. It maps the labels of all the clustering into corresponding binarized column vectors. The number of hyperedges, in this case, depends on the number of clusters. If the number of clusters is *k*^1^, *k*^2^, …, *k*^*p*^ then 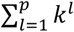 presents the total number of hyperedges. Here, *k*^*l*^ denotes the number of clusters in *l*^*th*^ clustering, and *p* denotes the number of clusterings. These indicator matrices can be collectively written, as *H* = (*H*^1^, *H*^2^, …, *H*^*p*^). It is also called the adjacency matrix of the hypergraph. Each of the column vectors of the hypergraph adjacency matrix is a specific hyperedge. Each row of the indicator matrix *H*^*l*^ represents the corresponding labels of clustering *l*. The key concept of the MCLA is to combine similar hyperedges and form meta-hyperedges. Later the instances (objects) are assigned to each of these meta-hyperedges based on the association membership values. Steps for the MCLA are the following:

1. Forming meta-graph from hyperedges
2. Transforming meta-graph into meta-clusters
3. Creating meta-hyperedges
4. Object association contest among meta-hyperedges

#### 2.3.4. Hybrid Bipartite Graph Formulation (HBGF)

The Hybrid Bipartite Graph Formulation treats clusters and data points as its basic entities. The first step in this algorithm is to create a bipartite graph and then partition the graph to obtain optimized clustering with optimal clusters. Each part of the partitioned bipartite graph represents the consensus cluster. For clustering (*C*_1_, *C*_2_, …, *C*_*n*_), a bipartite graph can be represented with *G*(*V, E*). Here, *V* represents the set of instances (*v*_1_, *v*_2_, …, *v*_*n*_), clustering (*C*_1_, *C*_2_, …, *C*_*n*_), and *E* represents the edges between the nodes. The edges are undirected, and every node in the graph has an edge with the other nodes. Each edge has a weight *W*(*i, j*) associated with it. Here, *i* and *j* represent the nodes. Edge weight between nodes *i* and *j* is defined as given in Equation 4.

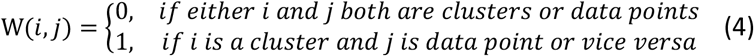

### 2.4. Evaluation Measures

#### 2.4.1. Test Accuracy and Subset Accuracy

To measure the node classification accuracy, one hot encoder is used as the label for all the nodes. Value 1 denotes the activation of the corresponding class, while value 0 depicts the deactivation of corresponding classes. The predicted outcome for a sample is called predicted matched only if both the outcome and label have one at the same place. If the total number of labels in the test set are *U*, and the total predicted matched sample outcomes are *V*, then the test accuracy (TA) is defined as 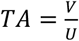. However, in the case of multi-label classification, a method such as a subset accuracy [55] can be used.

#### 2.4.2. Hamming loss and Hamming score

Hamming loss (HL) and Hamming score (HS) [56] were also used and they can be calculated as follow:

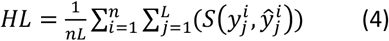

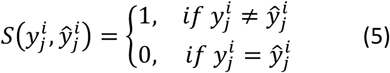

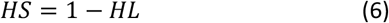

Here 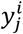 denotes the value of class *j* of label *i*, while 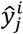 denotes the class *j* of the predicted outcome of sample *i*. L is the label size and *n* is the number of labels.

#### 2.4.3. Precision, Recall, and F-measure

The overlapping score between the ground truth complex *P* and predicted complex *Q* is defined as:

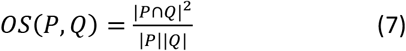

In this case the threshold for *OS*(*P, Q*) is set as 0.2 [57]. It denotes that if the value of the overlapping score between *P* and *Q* is 0:2, then both are matching each other. The Precision and Recall [58], [59] are defined as:

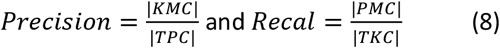

Where TKC is the Total Known Complexes, TPC is the Total Predicted Complexes, PMC is the number of Predicted Matched complexes, and KMC is the number of Known Matched Complexes. The F-Measure is calculated as follows:

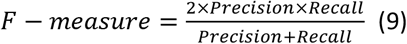

## 3. Experimental work and Results

### 3.1. Protein classification approaches

In this experimental work the Collins, Human, and Mouse datasets are used. The loss function was minimized through Adam optimizer with an initial learning rate of 0:01. The number of epochs is 200. The simple multi-class GCN classification model in which the length of each label is 2^*N*^ provided the test accuracy of 84.26% with 80:20 train and test data ratio on Collins dataset, where *N* is the total number of complexes (N = 10 for our experiment). The multi-label classification model provided the performance metrics with 100 communities, with Hamming Loss, Hamming Score, and Subset Accuracy of 0.0136, 0.9864, and 0.1554, respectively. To improve the performance, a modified version of the multi-class GCN method was used which offers greater flexibility in terms of the number of communities as well space and time complexities (please refer to section 2.2). In this case, the model was able to achieve 0.676 precision, 0.837 recall, and 0.748 F-score on Human dataset.

### 3.2. Representation Learning Approaches

In this experimental work, we aim to address the scarcity issue. The NOCD GCN model is implemented on 50 communities with the hidden size of 512, the weight decay is 1e-3, the learning rate is kept to 1e-4, the dropout rate is 0.05, the batch size is selected 20000. The number of epochs, in this case, was 200. The results of the NOCD GCN model on the Mouse dataset are presented in Table 2.

**Table 1:** Performance metrics of NOCD GCN model on Mouse dataset.

| Metrics | Value |
| --- | --- |
| NMI | 0.404 |
| Matched predicted complexes | 23 |
| Total predicted complexes | 50 |
| Matched known complexes | 30 |
| Precision | 0.60 |
| Recall | 0.46 |
| F-measure | 0.52 |

**Table 2:**
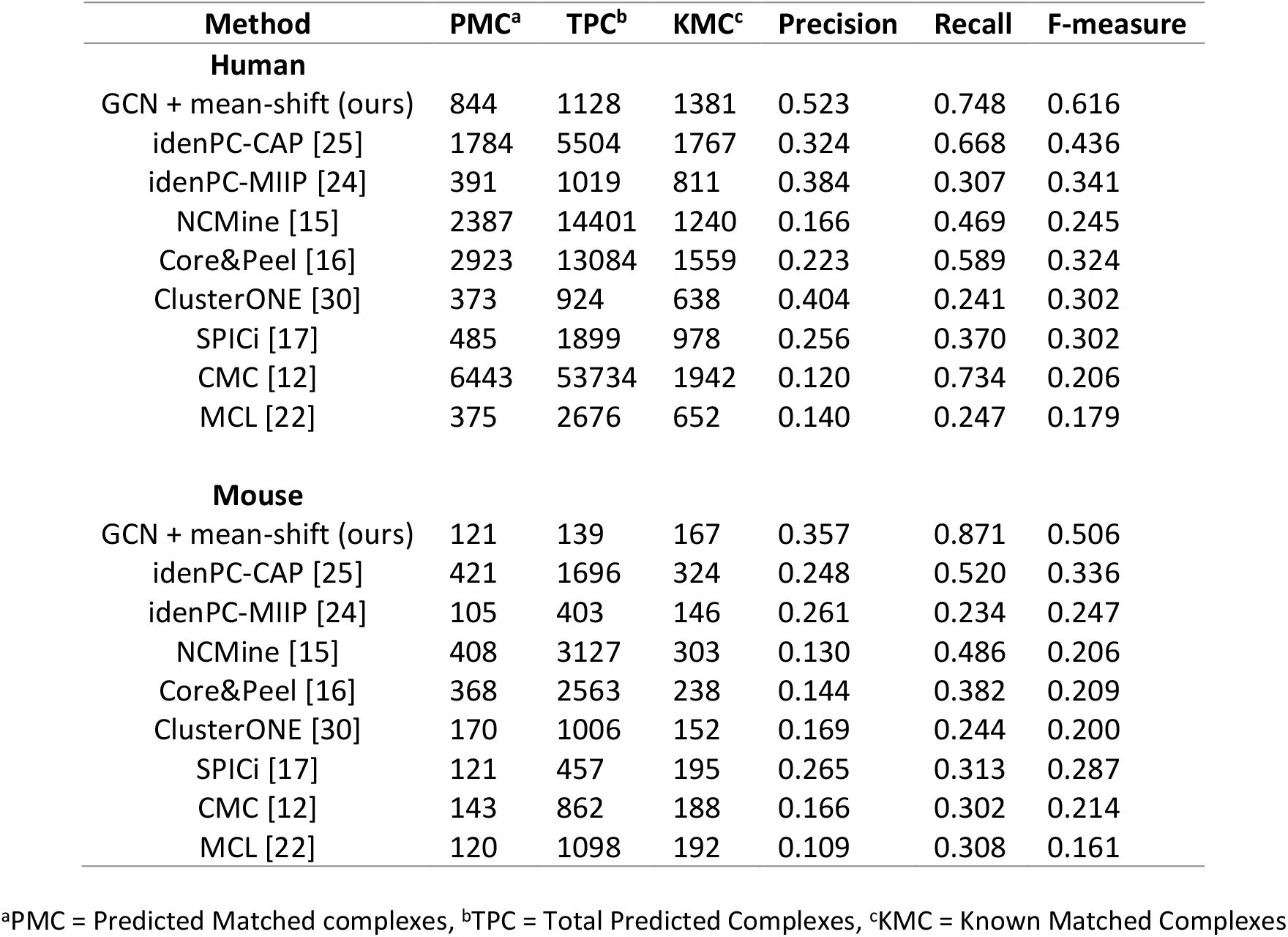
Overall performance comparison of (GCN + mean-shift) model with state-of-the-art methods on Human and Mouse datasets.

| Method | PMC<sup>a</sup> | TPC<sup>b</sup> | KMC<sup>c</sup> | Precision | Recall | F-measure |
| --- | --- | --- | --- | --- | --- | --- |
| **Human** |  |  |  |  |  |  |
| GCN + mean-shift (ours) | 844 | 1128 | 1381 | 0.523 | 0.748 | 0.616 |
| idenPC-CAP [25] | 1784 | 5504 | 1767 | 0.324 | 0.668 | 0.436 |
| idenPC-MIIP [24] | 391 | 1019 | 811 | 0.384 | 0.307 | 0.341 |
| NCMine [15] | 2387 | 14401 | 1240 | 0.166 | 0.469 | 0.245 |
| Core&Peel [16] | 2923 | 13084 | 1559 | 0.223 | 0.589 | 0.324 |
| ClusterONE [30] | 373 | 924 | 638 | 0.404 | 0.241 | 0.302 |
| SPICi [17] | 485 | 1899 | 978 | 0.256 | 0.370 | 0.302 |
| CMC [12] | 6443 | 53734 | 1942 | 0.120 | 0.734 | 0.206 |
| MCL [22] | 375 | 2676 | 652 | 0.140 | 0.247 | 0.179 |
| **Mouse** |  |  |  |  |  |  |
| GCN + mean-shift (ours) | 121 | 139 | 167 | 0.357 | 0.871 | 0.506 |
| idenPC-CAP [25] | 421 | 1696 | 324 | 0.248 | 0.520 | 0.336 |
| idenPC-MIIP [24] | 105 | 403 | 146 | 0.261 | 0.234 | 0.247 |
| NCMine [15] | 408 | 3127 | 303 | 0.130 | 0.486 | 0.206 |
| Core&Peel [16] | 368 | 2563 | 238 | 0.144 | 0.382 | 0.209 |
| ClusterONE [30] | 170 | 1006 | 152 | 0.169 | 0.244 | 0.200 |
| SPICi [17] | 121 | 457 | 195 | 0.265 | 0.313 | 0.287 |
| CMC [12] | 143 | 862 | 188 | 0.166 | 0.302 | 0.214 |
| MCL [22] | 120 | 1098 | 192 | 0.109 | 0.308 | 0.161 |

<sup>a</sup>PMC = Predicted Matched complexes, <sup>b</sup>TPC = Total Predicted Complexes, <sup>c</sup>KMC = Known Matched Complexes

As described in section 2.2, customized feature matrix has been constructed and NOCD GCN model has been applied using this feature matrix. On top 10 complexes of Human dataset, NMI score has been recorded as 0.5. NMI score started decreasing as we increase the number of complexes. For top 100 complexes, NOCD GCN method with customized feature matrix achieved NMI score of 0.105 on Human dataset.

Representation learning with the feature extracted through the pre-trained GCN from the Mouse and Human datasets has been performed with the clustering algorithm mean shift. The results of this feature learning algorithm are provided in Table 2.

The number of epochs, in this case, was 400 and the hidden sizes 512 are selected for this process. The remaining settings were kept the same as the multi-class GCN classification experimental work. It is quite clear to notice that the proposed approach for clustering proteins (nodes) outperformed all the state-of-the-art clustering algorithms. Hence, proving the effectiveness of the proposed approach.

Both MCLA and HBGF algorithms require a prior estimation of the number of clusters. Thus, we performed a grid search using the HBGF and MCLA to get the optimal number of clusters. Gavin dataset is used in this case to test this approach. First, three classical protein complex detection techniques (CMC [12], ClusterONE [30], and PEWCC [14]) were used to detect clusters in Gavin PPI dataset. The number of clusters yielded using the three techniques were 124, 243, and 206, respectively. After applying the MCLA and HBGF algorithms on the detected clusters using grid search from 60 complexes to 250 complexes. The best performances (MCLA method with 237 complexes and HBGF method with 66 complexes) are shown in Table 3.

**Table 3:**
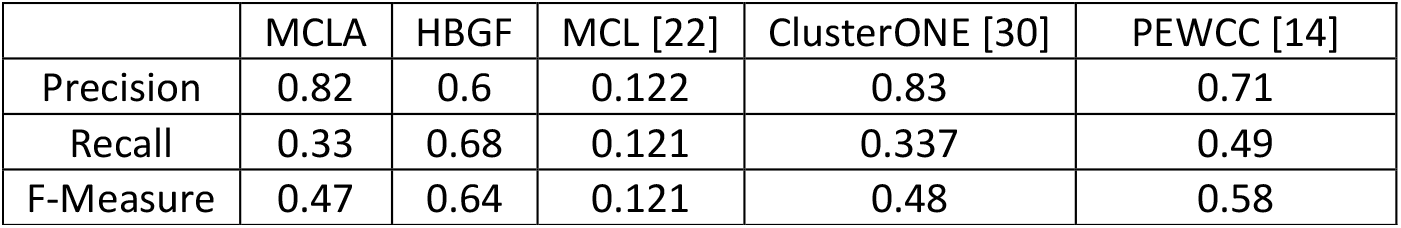
Performance metrics using MCLA and HBGF models

## 4. Discussion and Conclusion

In this paper, we introduced two GCN-based approaches to detect protein complexes in several benchmarked PPI datasets. The first approach is a multi-class GCN classification method and the second is a multi-label GCN classification method. The proposed approaches have overcome the detection accuracy and inefficiency in comparison to existing graph and density-based clustering approaches. We have also incrementally shown the improvements by changing the GCN method from a simple multi-class to multi-label classification problem and then improvised the multi-class GCN method.

Following the incremental improvements, we have proposed a sparse matrix operations-based GCN methodology. The efficiency of the time complexity in operations between the sparse and the dense matrices has also been compared. Then, we solved the problem of complex detection in an unsupervised manner. The NOCD model and (GCN feature extractor + mean shift clustering) method have been proposed. The performance of both approaches has been evaluated. The effectiveness of the proposed representation learning approach has been demonstrated on Human and Mouse datasets, in which it is shown to outperform the state-of-the-art methods when applied to detect the protein complexes. We have also shown the effect of the inclusion of explicit node features to the NOCD GCN method. Furthermore, assembling yielded clusters (complexes) by using the MCLA and HBGF algorithms has shown great potential. In the future, more complex detection methods can be added to the three shown in this experimental work to improve the precision and recall scores further.

Besides leveraging the advantages of the GCN, the proposed approaches were able to detect small complexes in which most of the state-of-the-art methods struggled to detect (as shown in Figure 3). For example, the proposed approaches accurately detected human complexes such as EXT1/EXT2 complex [60] [61], SUFU/GLIS3 complex, and NELF complex.

**Figure 3:**
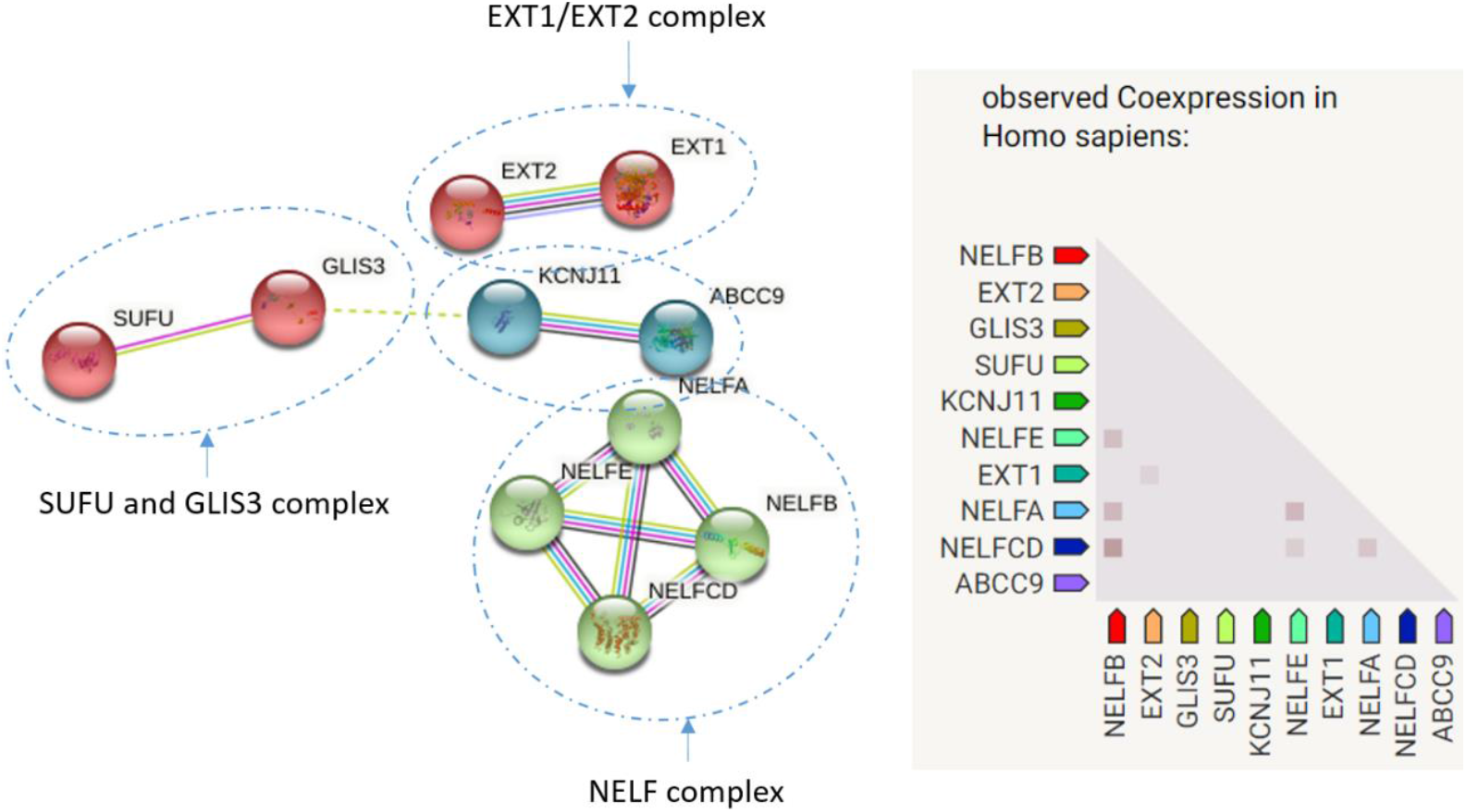
Samples of small human complexes detected 100% accurately by the proposed approaches which are hard to be detected by other methods.

Moreover, in this study, we used an identity matrix and a small length of customized feature vectors for providing node features. The length of the feature embedding can be further increased with the help of the topological features present in the neighborhood of the protein nodes. It forms the robust feature matrix for GCN operation, which might yield even better performance metrics.

A thorough investigation of the feature clustering algorithm has not been done in this research work. Mean shift algorithm has been selected from the pool of algorithms, which includes the Optics and affinity propagation method. Other techniques could also be tested and they may perform even better than the mean-shift algorithm.

## Glossary

GCN: Graph Convolutional Network
PPI: Protein-Protein Interaction
MCLA: Meta-Clustering Algorithm
NOCD: Neural Overlapping Community Detection
HBGF: Hybrid Bipartite Graph Formulation
PSO: Particle Swarm Optimization
OS: Overlapping Score
NMI: Normalized Mutual Information
RNA: Ribonucleic Acid
TKC: Total Known Complexes
TPC: Total Predicted Complexes
PMC: Predicted Matched complexes
KMC: Known Matched Complexes

## Acknowledgment

The authors would like to acknowledge partial support from the Big Data Analytics Center (BIDAC), United Arab Emirates University (UAEU).

